# Microsporidian coinfection reduces fitness of a fungal pathogen due to rapid host mortality

**DOI:** 10.1101/2024.02.08.579564

**Authors:** Marcin K. Dziuba, Kristina M. McIntire, Elizabeth S. Davenport, Emma Baird, Cristian Huerta, Riley Jaye, Fiona Corcoran, Paige McCreadie, Taleah Nelson, Meghan A. Duffy

## Abstract

Infection outcomes can be strongly context-dependent, shifting a host-symbiont relationship along a parasitism-mutualism continuum. Numerous studies show that under stressful conditions, symbionts that are typically mutualistic can become parasitic. The reverse possibility – a parasite becoming mutualistic – has received much less study. We investigated whether the parasitic microsporidium *Ordospora pajunii* can become beneficial for its host *Daphnia dentifera* in the presence of the more virulent fungal pathogen *Metschnikowia bicuspidata*. We found that, even though infection with *O. pajunii* reduces the frequency of penetration of *M. bicuspidata* spores into the host body cavity, it does not improve the survival or reproduction of the host; conversely, coinfection increased the mortality of *Daphnia*. However, the shorter lifespan of coinfected hosts disrupted the life cycle of *M. bicuspidata*, greatly reducing its fitness. Thus, coinfection with both pathogens was detrimental to the host at the individual level, but might be beneficial for the host population as a result of greatly reduced production of *M. bicuspidata* spores. If so, this would mean that *O. pajunii* outbreaks should delay or prevent *M. bicuspidata* outbreaks. In support of this, in an analysis of dynamics of naturally occurring outbreaks in two lakes where these pathogens co-occur, we found a time lag in occurrence between *O. pajunii* and *M. bicuspidata*, with *M. bicuspidata* epidemics only occurring after the collapse of *O. pajunii* epidemics. Thus, these results suggest that the interaction between co-occurring symbionts – and the net impact of a symbiont on a host – might be qualitatively different at different scales.

**Importance:** Understanding the factors that modify infection probability and virulence is crucial for identifying the drivers of infection outbreaks and modeling disease epidemic progression, and increases our ability to control diseases and reduce the harm they cause. One factor that can strongly influence infection probability and virulence is the presence of other pathogens. However, while coexposures and coinfections are incredibly common, we still have only a limited understanding of how pathogen interactions alter infection outcomes, or whether their impacts are scale-dependent. We used a system of one host and two pathogens to show that sequential coinfection can have a tremendous impact on the host and on the infecting pathogens, and that the outcome of (co-)infection can be negative or positive depending on the focal organization level.

## Introduction

We increasingly realize that whether a symbiosis is beneficial or harmful to the host can be context dependent and can rapidly shift. Those shifts occur due to changes in the costs and/or benefits of the interaction and can be triggered by changes in the environment (e.g., resource availability, change in abiotic parameters) or interactions with other organisms. For instance, mycorrhizal fungi are generally beneficial to their host plants, but the costs of symbiosis rise above the benefits when nutrients become abundant (1). In addition, female *Aedes albopictus* mosquitoes infected with *Wolbachia* have improved survival rates at low population densities, but are weaker competitors at high population densities (2). To date, most studies of shifts along a mutualism-parasitism gradient on ecological timescales have focused on organisms that are typically mutualists but that can become parasitic. Here, we consider the opposite: whether a symbiont that is parasitic can become a mutualist. More specifically, given that there are defensive mutualists that protect their hosts against pathogens (3–5), we consider the possibility that a symbiont that is typically parasitic might become a mutualist if it helps protect against a more virulent pathogen.

While scientists have generally overlooked the possibility that a pathogen might shift on ecological timescales from a pathogen to a mutualist based on protecting the host from a more virulent pathogen, it is well established that the presence of other microbes can alter the net outcome of interactions between hosts and their microbial symbionts. Multiple infections (or coinfections) are common in natural populations and the symbionts frequently interact with each other in diverse ways (6, 7). Exploitation of limited resources provided by the host can lead to strong competition between coinfecting pathogens, resulting in antagonism between the coinfecting symbionts or even competitive exclusion; on the other hand the coinfection can compromise the host’s immune system strongly enough to result in mutual facilitation of the pathogens (8).

In many cases, coinfecting pathogens arrive sequentially, with one arriving after the other is already established. In such cases of sequential infection, the fitness of the second arriving pathogen can be strongly impacted by the presence of the first. Often the expectation is that the fitness of the later arriving pathogen will be reduced, and there is, indeed, evidence for that. For example, initial infection of *Daphnia magna* hosts by a less virulent strain of the bacterium *Pasteuria ramosa* resulted in lowered fitness of a subsequently infecting highly virulent strain of *P. ramosa* (9). However, the late-arriving pathogen can also benefit from its host being formerly exposed to a different pathogen, as is the case for Israeli Acute Paralysis Virus, which benefited from its bumble bee host’s prior exposure to microsporidian *Nosema bombi* (10). The outcome of coinfection depends on the characteristics of the symbionts and the timing/order of infection (8, 11), and it can be strong enough to drive the evolution of virulence of the symbionts (12, 13).

In this study we investigate a recently discovered symbiotic relationship between a crustacean host, *Daphnia spp.,* and a microsporidium, *Ordospora pajunii* (14, 15). The microsporidium is parasitic by nature (14), infecting gut epithelial cells in its host, which then lyse as microsporidian spores are released back into the gut. However, in natural lakes, *Daphnia dentifera* was found to benefit from the infection when more virulent parasites were present in the environment. Specifically, *Daphnia* infected with *O. pajunii* produced more offspring than uninfected *Daphnia* during epidemics of obligate killer parasites like *Metschnikowia bicuspidata*, *Pasteuria ramosa,* or *Spirobacillus cienkowskii*; additionally, the guts of field-collected hosts infected with *O. pajunii* were less penetrable to *M. bicuspidata* spores (16), which infect by puncturing the gut epithelial cells. This suggests the possibility of a shift in the outcome of symbiosis contingent upon coexposure to a second pathogen.

In this study, we explored (a) how infection with *O. pajunii* affects the outcome of subsequent exposure to the strongly virulent fungus *M. bicuspidata* in controlled laboratory settings, (b) how interactions between the two symbionts affect the host and (c) whether these interactions might influence *M. bicuspidata* epidemics. To explore these questions, we sequentially exposed *Daphnia dentifera* to the microsporidian *O. pajunii* and fungal parasite *M. bicuspidata* in the lab, expecting to find (based on the previous field study (16)) evidence of a protective impact of *O. pajunii* infection against the fungus. We compared the number of fungal spores attacking guts of the hosts previously exposed or unexposed to the microsporidian. We looked at the lifespan, reproduction, fungal infection probability and spore burden in both host groups. We carried out this experiment in *ad libitum* and limited food conditions, expecting that the hypothesized beneficial effects of *O. pajunii* infection might diminish under limited resource availability; in the prior field study, the fitness impact of *O. pajunii* varied with resource levels (16). The results of our lab studies suggested that it might be difficult for *M. bicuspidata* to coexist with *O. pajunii* at the population-level. Therefore, we also analyzed data on prevalence of both symbionts in natural *D. dentifera* populations, expecting a temporal mismatch in their occurrence, potentially driven by the protective effects of *O. pajunii* against *M. bicuspidata* infections in *D. dentifera* populations.

## Methods

### Study system

This study focused on interactions between the host *D. dentifera* and two symbionts, *O. pajunii* and *M. bicuspidata*. *Daphnia*, a planktonic microcrustacean, is a model organism commonly used in ecology and evolutionary biology (17, 18). *O. pajunii* is a newly characterized gut-infecting microsporidium (14, 15) that is generally thought to be parasitic to *D. dentifera*, but can become beneficial in some circumstances, providing an interesting and rare example of ecologically driven shifts between parasitism and mutualism (16). The within-generation virulence of *O. pajunii* is rather weak: the most susceptible host genotypes suffer shortened lifespan and reduced fecundity, and less susceptible clones carry no detectable costs of infection (14). The spores of this microsporidian are continuously shed by the infected host (14). In contrast, *M. bicuspidata* is a highly virulent parasite; this fungus kills hosts relatively quickly and also reduces fecundity (19). It is an obligate killer, requiring host death for the parasite spores to be released into the environment.

### Experimental setup

The experiment was designed to test how prior infection with *O. pajunii* affects the probability of infection and virulence of *M. bicuspidata* under sufficient and scarce food resources. Therefore, we prepared two pathogen treatments: in the MB + OP treatment we sequentially exposed *D. dentifera* to *O. pajunii* on day 3 and then to *M. bicuspidata* on day 20; in the MB treatment we exposed *D. dentifera* just to *M. bicuspidata* on day 20 (with no exposure to the microsporidium). We also prepared control treatment that was not exposed to any parasites; in these, we dosed the *D. dentifera* with a placebo solution at each exposure day (placebo preparation described below), then monitored for background mortality. This treatment is not analyzed in this manuscript as it is not relevant to the hypotheses tested, but it demonstrates that animals were able to survive and reproduce under the experimental conditions when they were not exposed to a pathogen: Control high food (mean±SE) lifespan = 40±4.0 days and lifetime reproduction = 143±19.5 offspring; Control low food lifespan = 25±0.7 days and lifetime reproduction = 27±1.4 offspring.

The pathogen treatments were applied in *ad libitum* (High) and scarce (Low) resource levels, receiving daily concentrations of 20,000 or 1,000 cells/mL of the nutritious green algae *Ankistrodesmus falcatus*, respectively. These food concentrations were chosen based on a pilot experiment that indicated that reproductive output has saturated at the resource level in the high food treatment, while the low food treatment significantly reduces the reproduction of *Daphnia* (Supplementary Materials Figure S1). Each treatment had 30 replicates, resulting in 2 pathogen treatments × 2 food treatments × 30 replicates = 120 experimental units; however, some of the animals died before the day 20 exposure to MB, and hence were removed from the analyses as explained more below. The experiment began with third clutch offspring of parthenogenetically reproducing *D. dentifera*; we used the “S” genotype (referred to as “Standard” or “Std” in other publications on this system) because it is well-studied and highly susceptible to both of these pathogens. *D. dentifera* neonates were collected within 24h of birth, placed singly in 150 mL beakers filled with 100 mL of filtered lake water (collected from North Lake, Washtenaw County, Michigan, USA and filtered with Pall AE glass microfiber filters), randomly assigned to their respective pathogen (MB or OP+MB) and food (High or Low) treatments (day 0), and fed daily. On the third day, the water in all experimental beakers was replaced and the volume reduced to 50 mL, and OP+MB treatments were exposed to *O. pajunii* spores. The spores were prepared by harvesting infected animals from our *O. pajunii* ‘farm’ (an isolate of *O. pajunii* that was collected in Walsh Lake, Washtenaw County, Michigan, USA, and cultured in the S clone of *D. dentifera*, for details see 14); these animals were then combined and ground by hand in a single centrifuge tube using a plastic pestle. Each beaker in the OP+MB treatment then received a dose that was equivalent to that from a single donor host. At the same time, the MB treatment was dosed with a placebo consisting of uninfected *Daphnia*, with each beaker again receiving a dose equivalent to a single host. After 48h of exposure, each animal was moved to a new beaker filled with 100 mL of fresh filtered lake water. On day 20, after making sure that *O. pajunii* infections had fully developed in animals in the OP+MB treatment (by visual inspection under a stereomicroscope), all experimental animals were once more moved to 50 mL of filtered lake water and exposed to *M. bicuspidata* spores (250 spores/mL). We prepared the *M. bicuspidata* spore slurry by harvesting animals infected with *M. bicuspidata* from our farm (‘Standard’ isolate of *M. bicuspidata,* which is also cultured in the ‘S’ clone of *D. dentifera*) and grinding them using a motorized pestle; we estimated the spore density under the microscope with x400 magnification and using a hemocytometer, and then we dosed all animals with *M. bicuspidata* spores. The exposure lasted 48h, after which animals were placed in new beakers with 100 mL of fresh filtered lake water. At the end of *M. bicuspidata* exposure, hosts were inspected under the microscope with x200 magnification and attacking spores of *M. bicuspidata* (i.e., spores embedded in or penetrated through host’s gut (see 20, 21) were counted. Outside of the exposure periods, the water and beakers were changed once a week. Throughout the experiment, animals were kept at 20 ℃ and 16:8 light:dark. Animals were inspected daily for death and reproduction, and offspring were counted and removed from beakers. After death, animals were stored at -20℃; each animal that had lived long enough to survive exposure to *M. bicuspidata* (i.e. >22 days) was ground up and the abundance of *M. bicuspidata* spores was quantified using a hemocytometer and a microscope with x400 magnification. Defining infections based on the visible symptoms of *O. pajunii* infection prior to *M. biscuspidata* exposure, and by the presence of *O. pajunii* spores in the hemocytometer samples, the prevalence (successful infection) of *O. pajunii* reached 100% (30/30) in high food and 88% (23/26) in low food OP+MB individuals. Our analyses below included the 12% of animals from the low food OP+MB treatment that did not show these signs of infection, yielding a conservative measure of the effect of OP on MB.

### Field study

We sampled 15 lakes in Southeastern Michigan, USA in 2021 and 2022, and found *M. bicuspidata* and *O. pajunii* coexisting in two of them (Sullivan and Walsh Lakes in Washtenaw County, Michigan, USA). The lakes were sampled every two weeks from July to November, using a Wisconsin net (12 cm diameter, 153 µm mesh) that was vertically hauled from the bottom to the surface of the lake’s deepest spot. Each sample comprised three hauls, taken from spots located at least 5m apart. The samples were screened for density and infection status of at least 200 *D. dentifera* (or all individuals in the sample if it contained <200 *D. dentifera*, which follows established field protocols, e.g., 22, 23), using a dissecting microscope with dark field, and x10-115 magnification.

### Statistical analysis

We hypothesized that *O. pajunii* has a protective impact on its host when those hosts are exposed to *M. bicuspidata*, and that resource levels would also influence these interactions. Therefore, we looked at whether the number of *M. bicuspidata* spores attacking the guts of experimental *Daphnia* varied depending on the presence of *O. pajunii* and the food supply. We analyzed the data using a linear model with pathogen treatment and food level as explanatory variables and square-rooted number of attacking spores as the response variable; the transformation was performed after visual inspection of the distribution of residuals, as it improved homogeneity of variances and normality of the distribution of residuals. We compared the survival of *Daphnia* exposed to just *M. bicuspidata* to the animals sequentially exposed to the fungus and the microsporidian separately for each food treatment using Cox Proportional-Hazards models. We did not use a model combining both variables (i.e. pathogen treatment and food level) and their interaction, as the full model diagnostics indicated that the interaction term violated the assumptions of proportional hazards. A visual inspection of the data did not suggest an interaction, and an additive model (excluding the interaction) yielded results qualitatively consistent with the analysis done separately on each food level (i.e., significant effect of the pathogen treatment, but no effect of food level).

To analyze the effect of our treatments on reproduction of *D. dentifera*, we counted offspring produced over the entire lifespan and summed it to obtain their lifetime offspring production. We analyzed these data with a linear model, testing the impact of pathogen treatment, food level, and their interaction.

To further explore the potential protective effects of *O. pajunii* infection against *M. bicuspidata,* we analyzed whether the host’s initial infection with the microsporidian changed the fitness of the fungus, looking at infectivity (infection prevalence) and reproduction (spore burden) of the fungus. We analyzed the *M. bicuspidata* infection prevalence in two ways: i) in the first analysis, we designated all animals that had any *M. bicuspidata* spores, regardless of if the spores were mature or not, as infected and estimated the *M. bicuspidata* infection probability, ii) in the second analysis, only the animals who yielded mature (i.e., infectious) spores were regarded as effectively infected, and we estimated the *M. bicuspidata* effective infection probability. In both analyses, we used a generalized linear model with a binomial distribution (infected = 1, uninfected = 0), and pathogen treatment and food level as explanatory variables, and included their interaction in the model. None of the animals became infected in the low food OP+MB group, which made it impossible to analyze the contrasts between the groups in the full model. Instead, we ran two additional analyses: i) a comparison of high food MB vs. high food OP+MB and ii) a comparison of high food MB vs. low food MB. To analyze the spore burden, we used only individuals who were infected with *M. bicuspidata*. Due to violations of parametric test assumptions, we analyzed the data with a non-parametric Aligned Rank Analysis of Variance (24, 25), with the number of mature *M. bicuspidata* spores as the response variable and pathogen treatment and food level as explanatory variables, and included their interaction in the analysis.

For all the analyses of experimental data, we excluded the animals who died before or at day 20 (i.e., the day of exposure to *M. bicuspidata*). Moreover, for mortality analysis, we also removed two individuals in the High Food OP+MB treatment who did not become infected with *M. bicuspidata* and did not suffer from early mortality following the *M. bicuspidata* exposure (those animals lived 57 and 73 days). We removed these animals because our aim was to study the joint effect of *O. pajunii* and *M. bicuspidata* infection. The fact that two animals infected with *O. pajunii* avoided *M. bicuspidata*-driven early mortality (as opposed to none in the case of *D. dentifera* exposed just to the fungus) might indicate a protective function of *O. pajunii*, but just two cases are not enough to reliably analyze and draw any conclusions. The final sample sizes for the treatments in mortality analysis were: High Food MB: 24, High Food OP+MB: 27, Low Food MB: 23, Low Food OP+MB: 26.

We used the data on disease outbreaks in natural *D. dentifera* populations to look at whether *M. bicuspidata* and *O. pajunii* epidemics are temporally aligned (which could be a sign of facilitation) or if they are offset (potentially indicating antagonism between them) in the lakes in which they co-occur. We used cross-correlation analysis with *forecast* (26) and *zoo* (27) packages to estimate the time lag between the occurrence of the *O. pajunii* peak and *M. bicuspidata* peak in natural populations of *D. dentifera*.

All analyses were performed with R version 4.2.1 (28).

## Results

Consistent with a prior study of field-collected animals, *D. dentifera* infected with *O. pajunii* had fewer spores of *M. bicuspidata* attacking their guts in comparison to individuals without prior exposure to *O. pajunii* (mean attacking spores = 3.85 vs. 5.66; *F*_1,80_=6.008, *p*=0.016). More fungal spores attacked the guts of *Daphnia* in low food compared to high food in both pathogen treatment groups (mean attacking spores = 3.73 vs. 6.29; *F*_1,80_=8.319, *p*=0.005; Fig. 1); there was not a significant interaction between the pathogen and food treatments (*F*_1,80_=1.222, *p*=0.272). Overall, the hosts that were exposed only to *M. bicuspidata* in low food conditions had the most spores attacking their guts. Despite this, individuals in this treatment did not suffer the most from their interactions with *M. bicuspidata*. Instead, the mortality of individuals with prior exposure to *O. pajunii* was greater than that of animals unexposed to *O. pajunii* in both high food (hazard ratio=3.775, 95% CI 1.986 to 7.175, *p*<0.001) and low food (hazard ratio=26.991, 95% CI 7.48 to 97.4, *p*<0.001) (Fig. 2). This increased mortality coincided with lower reproduction of *Daphnia* exposed to *O. pajunii* (pathogen treatment effect on lifetime offspring production *F*_1,97_=44.338, *p*<0.001; Fig. 2); as expected, there was also a strong effect of food level on reproduction (food treatment effect *F*_1,97_=447.528, *p*<0.001; pathogen×food interaction *F*_1,97_=0.275, *p*=0.601; Fig. 2).

**Figure 1.**
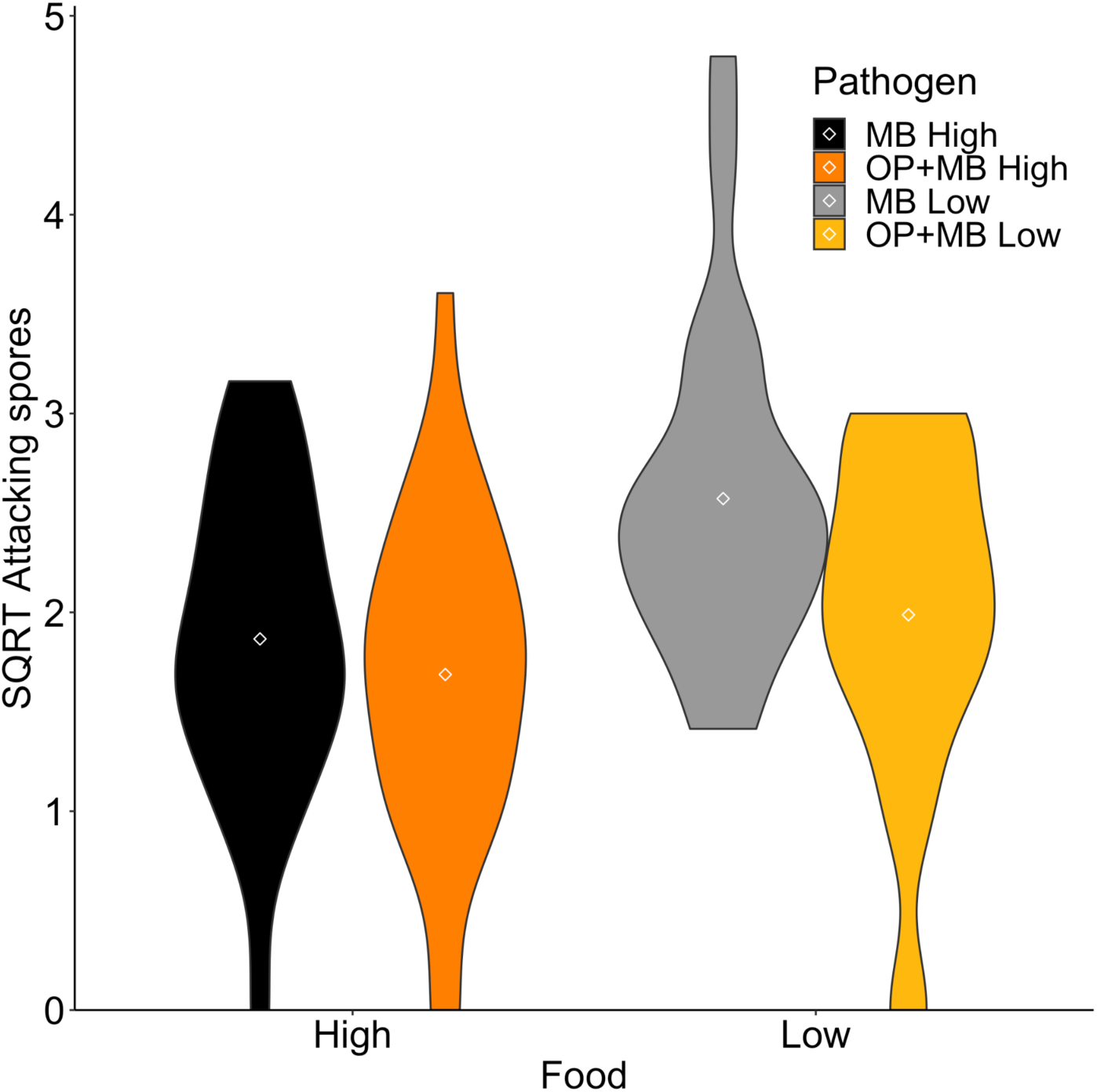
Fewer *M. bicuspidata* spores attacked the guts of *Daphnia* infected with *O. pajunii,* as compared to uninfected hosts (*F*_1,80_=6.008, *p*=0.016). In addition, hosts reared on lower food concentrations had more attacking spores (*F*_1,80_=8.319, *p*=0.005). There was not a significant interaction between pathogen treatment and food level (*F*_1,80_=1.222, *p*=0.272). ‘MB’ indicates hosts just exposed to *M. bicuspidata*, whereas ‘OP+MB’ indicates hosts exposed first to *O. pajunii* and then to *M. bicuspidata*.

**Figure 2.**
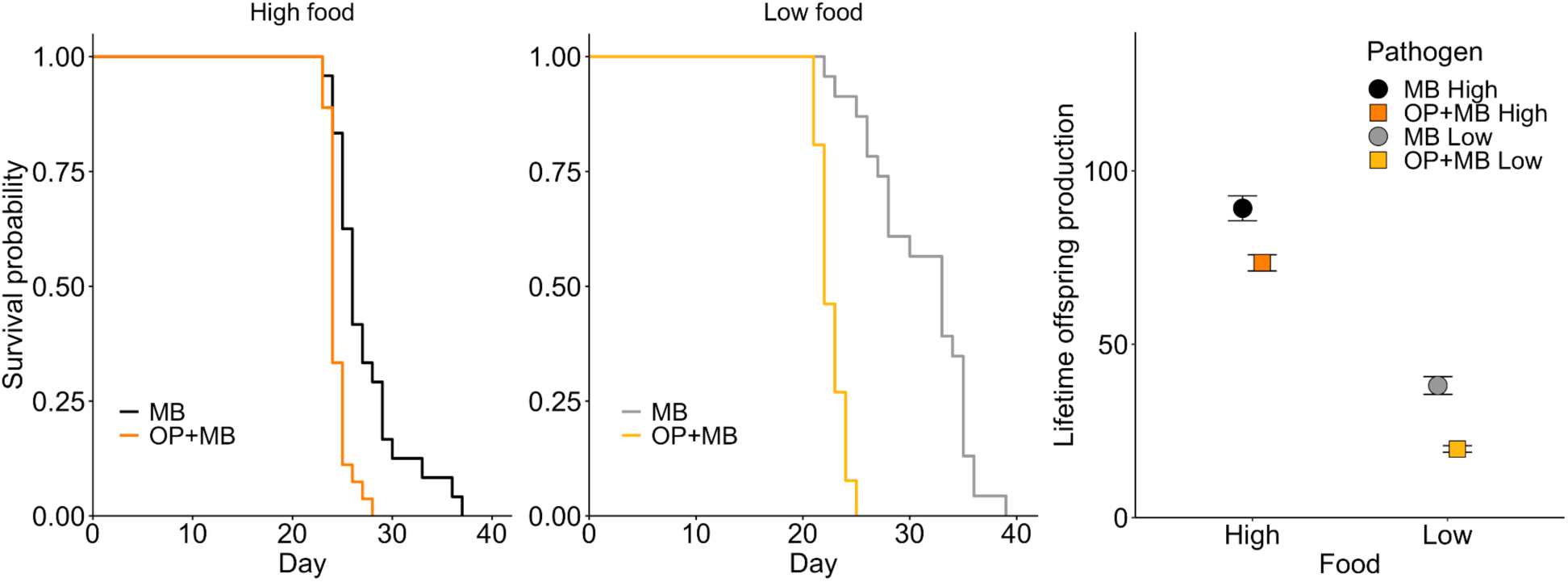
In both high and low food treatments, *D. dentifera* exposed to *M. bicuspidata* had a shorter lifespan when they were already infected with *O. pajunii* (left panel: high food hazard ratio=3.775, 95% CI 1.986 to 7.175, *p*<0.001; middle panel: low food hazard ratio=26.991, 95% CI 7.48 to 97.4, *p*<0.001). ‘MB’ indicates hosts just exposed to *M. bicuspidata*, whereas ‘OP+MB’ indicates hosts exposed first to *O. pajunii* and then to *M. bicuspidata.* Right panel: *Daphnia* infected with *O. pajunii* and exposed to *M. bicuspidata* produced fewer offspring than *Daphnia* just exposed to *M. bicuspidata* in both food regimes (parasite effect *F*_1,97_=44.338, *p*<0.001). As expected, food level also strongly influenced reproduction (food treatment effect *F*_1,97_=447.528, *p*<0.001). There was not a significant interaction between pathogen treatment and food level (*F*_1,97_=0.275, *p*=0.601).

A critical question in our study was whether *O. pajunii* can protect its host from becoming infected with the more virulent pathogen *M. bicuspidata* and hence reduce the propagation of the latter pathogen. Pathogen treatment interacted with food level to affect the probability of infection (pathogen treatment × food level interaction *χ*^2^_1_=20.987, *p*<0.001). At high food levels, *D. dentifera* were equally likely to become infected with *M. bicuspidata* in both pathogen treatments (pathogen treatment effect *χ*^2^_1_=0.270, *p*=0.604); however, when food was scarce, animals previously exposed to *O. pajunii* did not become infected with *M. bicuspidata* (Fig. 3, left panel), while those without *O. pajunii* exposure suffered greater infection risk than their high food counterparts (food treatment effect *χ*^2^_1_=7.497, *p*=0.006). The increased mortality observed among animals exposed to *O. pajunii* (Fig. 2) shortened the time available for *M. bicuspidata* to form mature, infectious spores. Therefore, we analyzed the probability of infection that results in the production of mature (i.e., infectious) spores, which we refer to as ‘effective infection’. In the high food treatment, *D. dentifera* infected with *O. pajunii* were less likely to produce effective *M. bicuspidata* infections, in comparison to *Daphnia* unexposed to *O. pajunii* (pathogen treatment effect *χ*^2^_1_=4.394, *p*=0.036), and as previously stated, in the low food treatment the animals infected with *O. pajunii* did not become infected with *M. bicuspidata* (Fig. 3, middle panel). In contrast, *D. dentifera* not exposed to *O. pajunii* were more likely to become ‘effectively infected’ when the food level was low as compared to when it was high (food treatment effect *χ*^2^_1_=4.329, *p*=0.037, Fig. 3b). Additionally, the number of mature *M. bicuspidata* spores produced in infected animals was drastically lower when they were initially exposed to *O. pajunii* (pathogen treatment effect *F*_1,23_=4.787, *p*=0.039); *M. bicuspidata* spore burden was not affected by the food level (*F*_1,23_=1.439, *p*=0.242).

**Figure 3.**
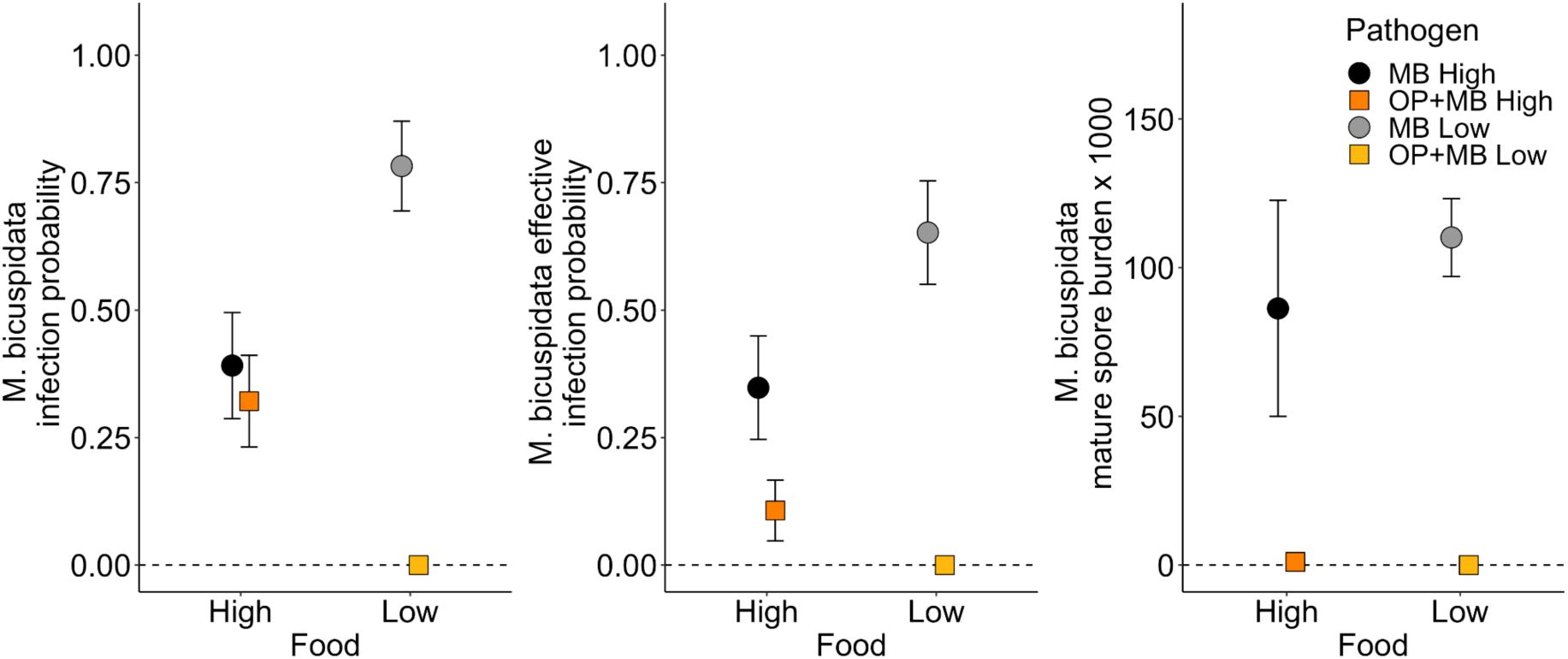
*Daphnia* infected with *O. pajunii* are as likely to become infected with *M. bicuspidata* as uninfected *Daphnia* under high food conditions; low food concentration reduces probability of infection with *M. bicuspidata* in *Daphnia* infected with *O. pajunii* to zero, but increases the probability of infection in *Daphnia* unexposed to *O. pajunii* (left panel). In other words, there is a strong interaction between *O. pajunii* infection and food level (*χ*^2^_1_=20.987, *p*<0.001), with *O. pajunii* protecting against *M. bicuspidata* infections in low food conditions. However, when looking just at the effective infections (i.e., those that result in production of viable transmission spores; middle panel), *Daphnia* infected with *O. pajunii* were much less likely to produce effective infections than uninfected *Daphnia* regardless of the food regime (parasite effect for high food treatment *χ*^2^_1_=4.394, *p*=0.036). In addition, even considering just those animals that developed effective *M. bicuspidata* infections, individuals that had been exposed to *O. pajunii* produced extremely few mature *M. bicuspidata* spores (right panel). Points and whiskers indicate mean and standard error.

Because our laboratory results suggested that infections with *O. pajunii* should greatly hinder propagation of *M. bicuspidata*, we looked at whether there was a temporal mismatch in the timing of disease outbreaks in lakes in which they co-occur. We found that *O. pajunii* and *M. bicuspidata* outbreaks were temporally offset, with *M. bicuspidata* reaching its peak about 6-8 weeks after the peak prevalence of *O. pajunii* (Fig. 4, Table 1); this lag was statistically significant in three of the four lake-years.

**Figure 4.**
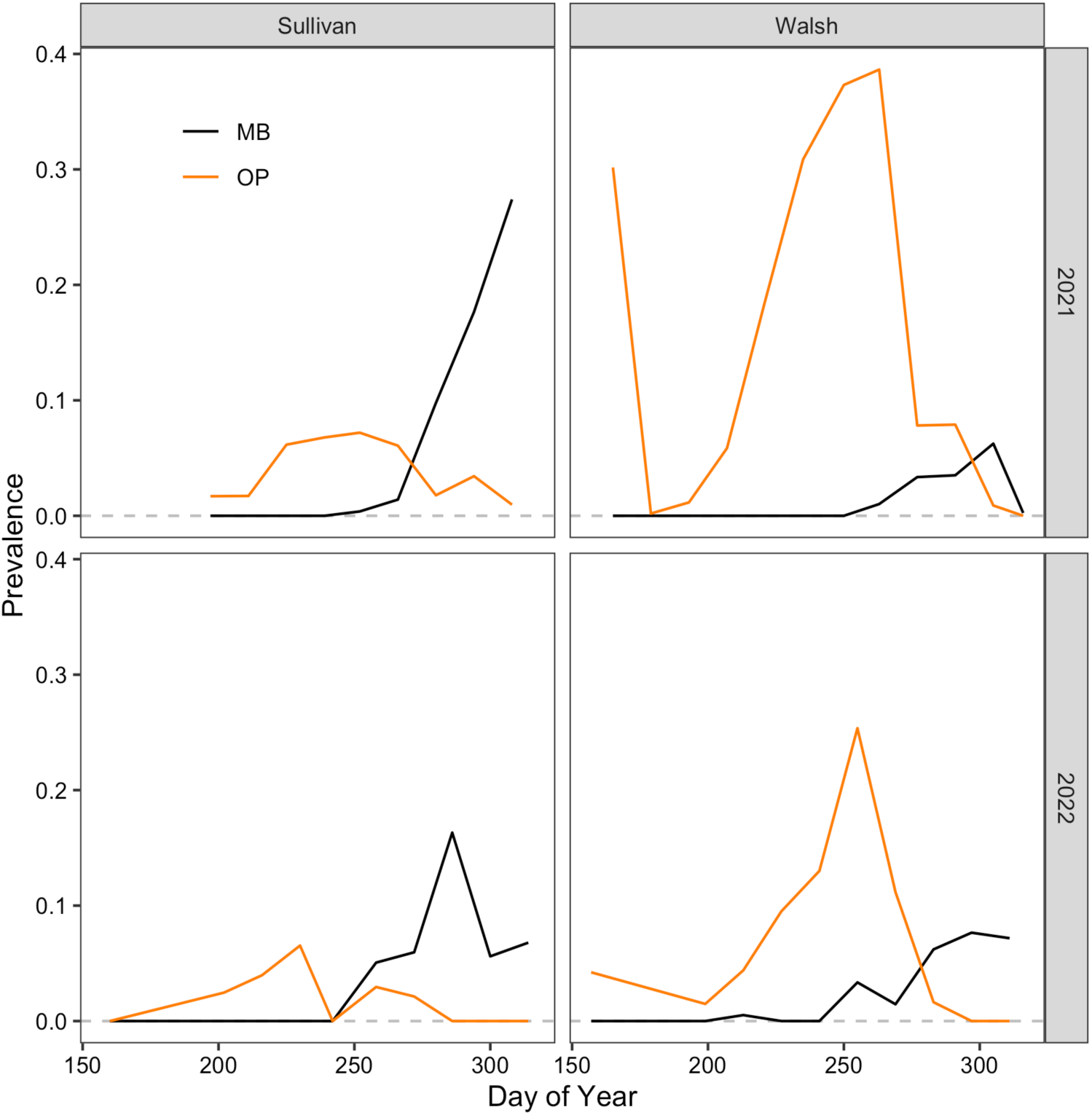
*M. bicuspidata* outbreaks in *D. dentifera* populations peak after *O. pajunii* prevalence decreases. Peaks of *O. pajunii* and *M. bicuspidata* were lagged by 6-8 weeks, with that lag being significant in three of the four lake-years (see also Table 1).

**Table 1.**
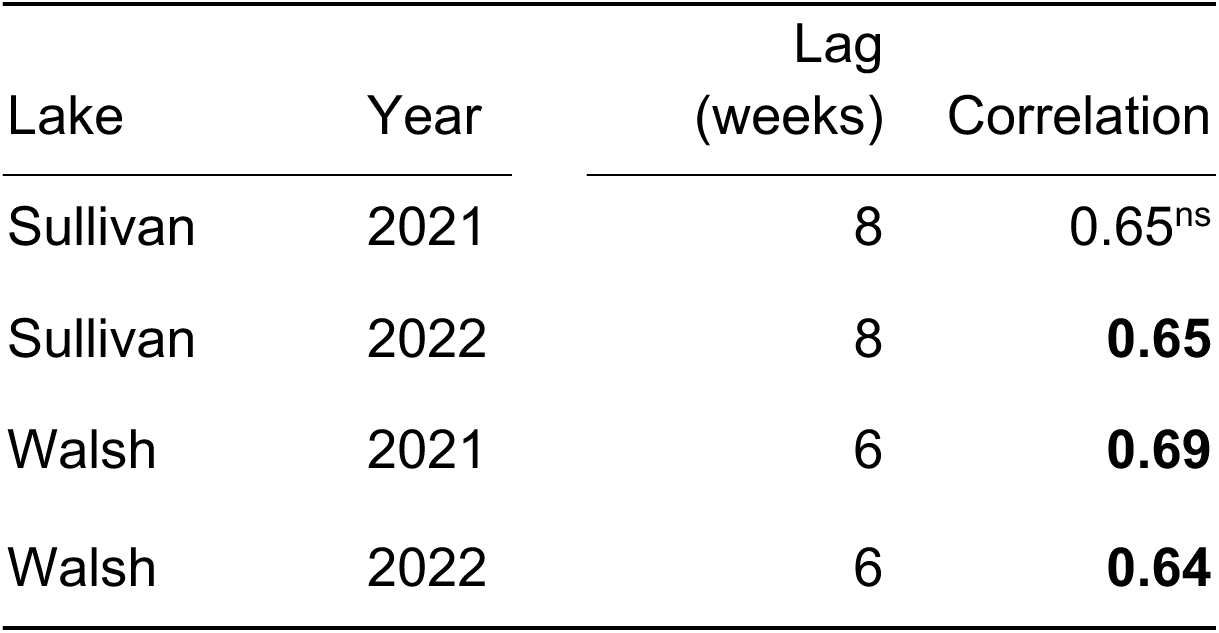
Lag between *O. pajunii* peak prevalence and *M. bicuspidata* prevalence in *D. dentifera* population, and correlation between the two symbionts at the lag. All the correlation coefficients are statistically significant except for Sullivan at 2021 (^ns^), which is marginally significant (i.e. the exact correlation coefficient is 0.646 and the significance margin (confidence interval) is at 0.653).

## Discussion

Our study explored whether a symbiont can shift from parasitism to mutualism depending upon the presence of another pathogen infecting the same host. We found that *O. pajunii* infections protected *D. dentifera* from attack by *M. bicuspidata* spores. This finding is in line with the previous results obtained with field collected *D. dentifera* (16). However, the effect size in our laboratory study was small, and *O. pajunii* infection did not improve the survival of *Daphnia* exposed to *M. bicuspidata*. Instead, *Daphnia* exposed to the fungus suffered high mortality, and prior infection with *O. pajunii* exacerbated the negative effect of fungal exposure/infection. Animals that were exposed to *O. pajunii* prior to *M. bicuspidata* had shorter lifespans and reduced lifetime reproduction. These results indicate that, from the perspective of an individual host, *O. pajunii* infection does not increase host fitness during *M. bicuspidata* exposure, even though it does protect against attacking spores; instead, sequential coinfection with *O. pajunii* and then *M. bicuspidata* was more harmful to the host than infection with only *M. bicuspidata* (or just *O. pajunii* (14)). Interestingly, the lifespan reduction of the host in the coinfection treatment narrowed the time window for the virulent parasite *M. bicuspidata* to develop, resulting in fewer *Daphnia* reaching the final stage of infection and a drastic reduction of the number of fungal spores produced in infected hosts. Thus, the reduction in host lifespan associated with coinfection resulted in a large reduction in fitness of the fungal parasite. Looking at these results from the host population perspective, *O. pajunii* could be beneficial for *Daphnia*, as it potentially reduces the size of, or even prevents, *M. bicuspidata* outbreaks.

Coinfection can result in increased virulence (e.g. 29, 30), especially when the pathogens compete for limited resources (12). Multiple infections often favor higher virulence, as more virulent parasites exploit their hosts more effectively, which allows them to increase their own reproduction and transmission (8, 12). However, our results show that high virulence in coinfections is detrimental to *M. bicuspidata*, as the rapid mortality of the host drastically reduces the probability of infection leading to subsequent transmission and the number of infectious spores produced by the fungus. *D. dentifera* hosts with established *O. pajunii* infections were of extremely low competence for *M. bicuspidata*; those animals that became infected with the fungus produced few, if any, mature spores. Similarly, a recent study found that infection with a congener of *O. pajunii* – *Ordospora colligata* – reduced the fitness of subsequently infecting *M. bicuspidata* due to higher mortality in the host *Daphnia magna* (31). Another study discovered that *O. colligata* reduces fitness of another microsporidium *Hamiltosporidium tvaerminnensis* upon coinfection, also because of increased host mortality (32). It seems unlikely that the strong virulence observed in our study was a result of resource depletion due to the symbionts, because *M. bicuspidata* had very little time to proliferate and exploit the host, and the coinfected group died earlier even when provisioned with high food. We consider it more likely that the damage caused by both pathogens was too severe for the host to withstand – *O. pajunii* infects the host’s gut epithelium and leads to cell breakdown at the end of the pathogen’s life cycle (14, 15), while needle-shaped spores of *M. bicuspidata* pierce through host’s gut, causing physical damage and triggering an immune response (21). However, at present this is speculative, and more extensive research is needed to establish the mechanism that shortens the lifespan of coinfected *Daphnia*.

*Ordospora pajunii* heavily constrained the reproduction of *M. bicuspidata*, which likely has a strong impact on the transmission rate of the latter. This in turn might have large ecological and evolutionary consequences. Outbreaks of pathogen epidemics in lakes frequently follow a rapid increase in the concentration of pathogen transmission stages in the water column (33). In the case of *M. bicuspidata*, outbreaks tend to occur in the fall, after the temperature reduction breaks thermal stratification and turnover enables pathogen spore resuspension from the sediment bank (34). This suggests that *M. bicuspidata* outbreaks are regulated by weather events (and solar radiation (35)). Our experiment indicates that if the host populations are already undergoing *O. pajunii* epidemics during potential *M. bicuspidata* outbreaks, the transmission potential of the latter becomes limited. Thus, *O. pajunii* might be another factor affecting *M. bicuspidata* population dynamics, shifting the timing of emergence, narrowing the time of epidemics, reducing infection prevalence or even preventing an outbreak. Our field data show that when the two pathogens coexist in the same lake, *M. bicuspidata* emergence is always preceded by *O. pajunii* epidemics, and that the peak of the fungal epidemics most often comes after *O. pajunii* prevalence declines. This is consistent with the emergence of *M. bicuspidata* being obstructed by the microsporidian (even if *M. bicuspidata* spores are in the water column, see 33, 36), with host individuals infected with *O. pajunii* that are subsequently exposed to and infected with *M. bicuspidata* dying before the latter can amplify and transmit. If that’s the case, *M. bicuspidata* would require a reduction of *O. pajunii* prevalence in order to emerge. It is interesting the *M. bicuspidata* suffers from not having enough time to develop during coinfections with *O. pajunii*, as prior work has shown that it can prevent a different parasite, the bacterium *P. ramosa*, from being able to complete its development within coinfected hosts (11).

Coinfection can be a strong selective factor, shaping the evolution of parasites. The outcome of selection is often difficult to predict, as it depends on multiple factors, including the nature of intra-host interactions, the host immune response, and whether the infection is simultaneous or sequential (for reviews see 8, 12). In many cases, coinfections should promote either highly virulent parasite genotypes or those capable of increasing virulence via phenotypic plasticity during coinfection (12). In the case of the *O. pajunii* and *M. bicuspidata* interaction, it seems that the addition of *O. pajunii* might drive selection on *M. bicuspidata.* One possibility is that it might select for more rapid growth by *M. bicuspidata*, so that it is able to complete development prior to host death. However, an experimental evolution study that attempted similar selection failed to produce a response in *M. bicuspidata* (37). Alternatively, if slower parasite growth reduces parasite virulence and hence host mortality (consistent with the idea that higher virulence is associated with less prudent resource use (38)), there could be selection for *slower* growth of *M. bicuspidata* in response to *O. pajunii*. Unfortunately, experimental tests of this in this system seem unlikely to be successful, as multiple attempts to select *M. bicuspidata* for a variety of conditions have failed (37, 39, 40). Instead, we propose that studies that quantify how the age and health state of the host (41, 42), the timing and order of exposures (11, 43) and the genotype of each organism (42, 44) influence outcomes during coinfections would likely be more fruitful.

This study was motivated by a general hypothesis that *O. pajunii* acts as a mutualist when its host is subsequently exposed to highly virulent *M. bicuspidata*. We found benefits (reduced number of attacking *M. bicupidata* spores) and costs (shorter lifespan, reduced reproduction) of *O. pajunii* infection. Whether the interaction is mutualistic or parasitic depends on the net outcome of costs and benefits accrued over the lifetime of the host (45). From the perspective of a single infected individual, *O. pajunii* is a parasite. However, the strong reduction of fitness of *M. bicuspidata* expressed during sequential coinfection with *O. pajunii* indicates that the across-generation effects might be non-trivial and potentially highly beneficial for the host. Given this, sacrificing a few members of the population slightly earlier (in the case of coinfection) than it would be if the host wasn’t infected with *O. pajunii* might be a small price for stopping the outbreak of more virulent pathogens, particularly in the case of a highly related population or clonally reproducing hosts, as in the case of *Daphnia*. This possibility falls into the category of a hypothesized phenomenon known as adaptive suicide – a mechanism of kin protection that has some empirical support as a response to parasites (46) and to competitors (47).

Whether *O. pajunii* could be considered a mutualist based on the population-level impacts of infection is controversial, as whether something is a mutualist or a parasite is traditionally defined based on the impact on the fitness of a host individual. However, our results suggest that an interaction that is harmful to a host individual might end up benefiting the population as a whole. Future studies that experimentally test this hypothesis at the population level – and that grapple with the potential for different impacts at different levels of biological organization – will help further unravel factors that drive shifts along a mutualism-parasitism continuum.

## Notes

### Competing Interest Statement

The authors have declared no competing interest.

