## Supplementary Materials for "Microsporidian coinfection reduces fitness of a fungal pathogen due to rapid host mortality"

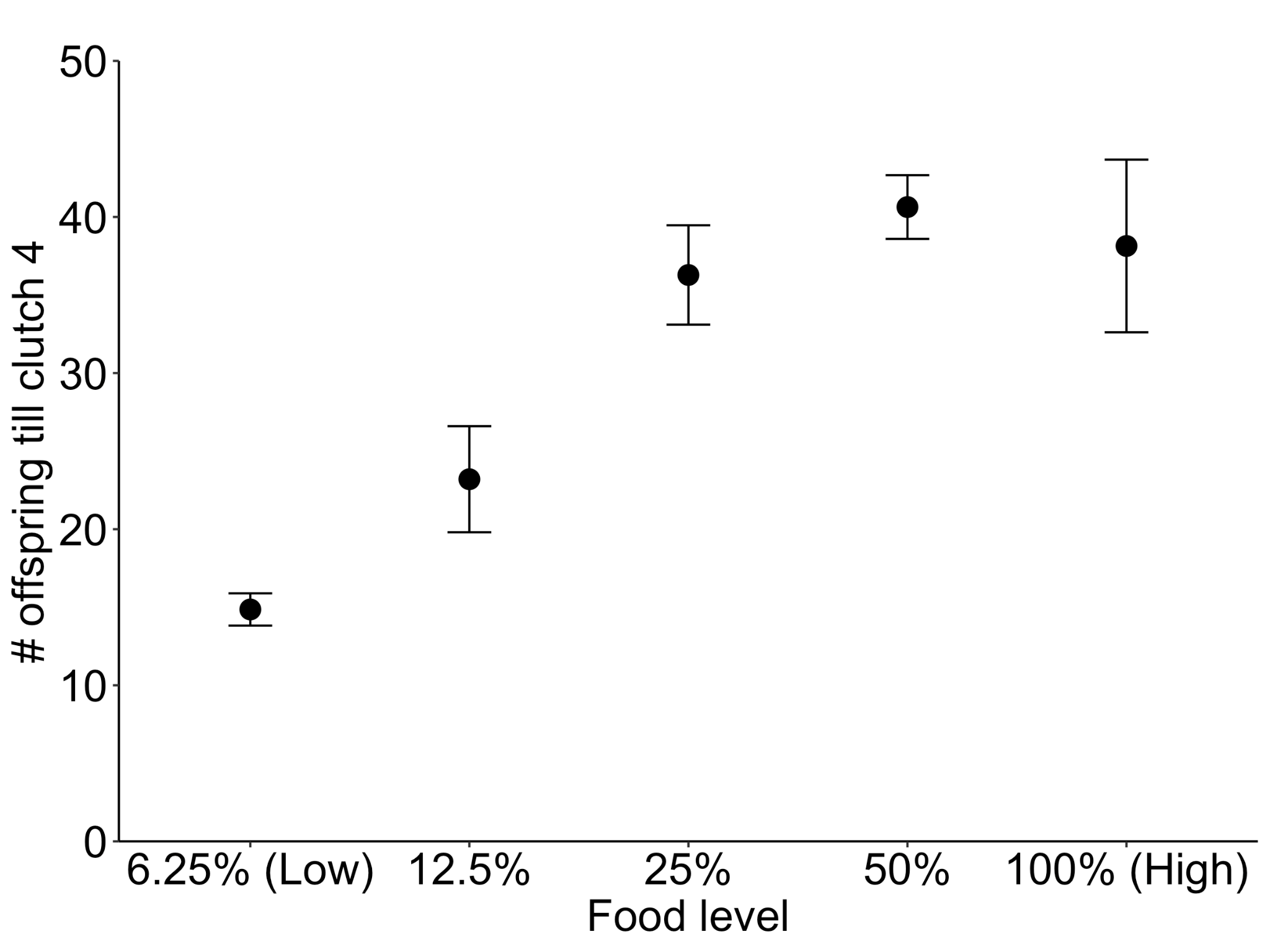

Figure S1. Total number of offspring produced during the first four clutches by uninfected *Daphnia dentifera* (clone “S”) in different concentrations of food, spanning from 1,000 cells/mL (6.25%) to 20,000 cells/mL (100%) of *Ankistrodesmus falcatus*. The highest and the lowest concentrations were selected as experimental high and low food concentrations, respectively, for the experiment in the main text. The highest concentration (100%) is a standard concentration fed to *Daphnia* as non-limiting. Dots and whiskers represent the averages and standard errors.
